# Dynamic copy number evolution of X- and Y-linked ampliconic genes in human populations

**DOI:** 10.1101/228841

**Authors:** Elise A. Lucotte, Laurits Skov, Moisès Coll Macià, Kasper Munch, Mikkel H. Schierup

## Abstract

Ampliconic genes are good candidates for speciation genes: they are testis-expressed, multicopy and localized on sex chromosomes. Moreover, copy number variation in a specific ampliconic gene pair (*Slx* and *Sly*) is involved in hybrid incompatibilities between *M. musculus* and *M. domesticus*. However, we know little about the distribution of the ampliconic genes copy number and their turnover in human populations. Here we explore the evolution of human X- and Y-linked ampliconic genes by investigating copy number variation (CNV) and coding variation between populations using the Simons Genome Diversity Project. We develop a method to assess CNVs using the read-depth on modified X and Y chromosome targets containing only one repetition of each ampliconic gene. Our results reveal extensive standing variation in copy number both within and between human populations for several ampliconic genes. For the Y chromosome, we can infer multiple independent amplifications and losses of these gene copies even within closely related Y haplogroups, that diversified less than 50,000 years ago. For the X chromosome, we also find high copy number and coding diversity within populations. While we cannot rule out that neutral processes are at the origin of this high diversity, this study gives insights on the distribution of copy number within human populations, and demonstrates an extremely fast turnover in copy number of these regions.

## INTRODUCTION

Ampliconic genes are only found on sex chromosomes and consist of several adjacent duplications of small genomic regions with more than 99.9% similarity between copies (Skaletsky et al. 2003). Their evolutionary turnover is very rapid: only 31% of human X ampliconic genes have an ortholog in mice compared to 95% for single copy genes (Mueller et al. 2013). Most of the human ampliconic genes are protein coding and expressed exclusively in the testis, however, their specific function in gametogenesis is poorly understood.

It is a long-standing hypothesis that intragenomic conflicts, *i.e.* an arms race between the X and the Y for transmission to the next generation, due to meiotic drive leads to an increased divergence between closely related species and therefore is at the origin of hybrid incompatibilities (Frank 1991; Hurst and Pomiankowski 1991). In mice, the homologous ampliconic genes *Slx* and *Sly*, located on the X and Y chromosome respectively, were co-amplified as a result of an intragenomic conflict (Soh et al. 2014). An unbalanced copy number of *Slx* and *Sly* leads to deleterious X-Y dosage disruption in hybrids (Larson et al. 2016) and a deficiency in *Slx* provokes a sex ratio distortion in offspring towards males, which is corrected with *Sly* deficiency (Cocquet et al. 2012). More generally, misregulation of the X and Y chromosome expression during spermatogenesis is pervasive in mice hybrids, affecting meiotic sex chromosome inactivation, a crucial phenomenon during male meiosis (Larson et al. 2016). While the X and Y-linked genes are repressed during meiosis, post-meiotically, genes in multiple copies have been shown to be expressed in round spermatids, dependent on their copy number (Mueller et al. 2008). Similar results were obtained in felids: fertility eQTL were mapped near X-linked ampliconic genes, and sterile hybrids show an overexpression of the X chromosome during meiosis compared to controls (Davis et al. 2015), suggesting that this phenomenon is widespread in mammals.

We recently reported that in primates, the X chromosome has mega-base wide losses of diversity, overlapping between species (Dutheil et al. 2015; Nam et al. 2015) and most probably resulting from recurrent selective sweeps. They overlap significantly with genomic areas depleted of Neanderthal introgression in humans, suggesting that the same regions are involved in creating hybrid incompatibilities. These regions are enriched for ampliconic genes, which prompted us to suggest that they may be speciation genes in primates (Nam et al. 2015).

While these observations point toward an important role of the ampliconic genes in speciation, little is known about the worldwide distribution of copy number variations in human populations of both X- and Y-linked ampliconic genes, as well as their dynamic of amplification.

Most of the recent studies on ampliconic genes copy number have focused on the Y chromosome, and were performed on one human population (62 Danes, Skov et al. 2017) or a few individuals from several primate species(Ghenu et al. 2016; Oetjens et al. 2016). To our knowledge, no population-wide studies on copy number variations of X-linked ampliconic genes have been conducted. However, a majority of them are part of the cancer/testis (CT) gene family and have been studied as targets for therapeutic cancer vaccines (see Simpson et al. 2005 for a review). Also, using comparative genomics between human and primate sequences, previous studies reported signals of diversifying selection in CT genes(Stevenson et al. 2007; Gjerstorff and Ditzel 2008; Liu et al. 2008; Zhao et al. 2012; Zhang and Su 2014) as well as recent amplification in the human lineage for GAGE (Gjerstorff and Ditzel 2008; Liu et al. 2008) and CTAGE families (Zhang and Su 2014).

Therefore, a worldwide-scale description of copy number variations in both X-and Y-linked ampliconic genes in human populations is lacking. Such investigation is critical to understand ampliconic gene evolution and their involvement in the emergence of hybrid incompatibilities. Indeed, if human ampliconic genes are involved in intragenomic conflicts and incompatibilities arise depending on their copy number, describing their copy number distribution and dynamic of amplification constitutes a first step in evaluating the importance of these genes in speciation and meiotic drive. Mueller et al. (2013) have improved considerably the human X chromosome reference genome for ampliconic genes (hg38), using single-haplotype sequencing, thus allowing us to investigate these regions at the population scale, as it was done in Danes for the Y-linked ampliconic genes (Skov et al. 2017).

Here, we investigate the evolutionary dynamics of ampliconic genes on the X and Y chromosomes in humans from the Simons Genome Diversity Project dataset (Mallick et al. 2016) which provides genomic sequencing for 128 human populations including 276 individuals (102 females, 174 males). We both surveyed ampliconic genes copy number variations and their coding sequence turnover. Due to their highly repetitive nature, classical methods could not be used, we therefore develop new bioinformatic approaches to assess their copy number and nucleotide variability. We find very dynamic evolution suggesting high mutation rates of these regions. While we cannot disentangle neutral processes from diversifying selection, this study provides the first global picture of the diversity and turnover of the ampliconic genes in human populations at a world-wide scale.

## MATERIALS AND METHODS

### Identifying the unit of repetition of each ampliconic regions

#### X chromosome

See figure 5 for a schema the method.

**Figure 5.**
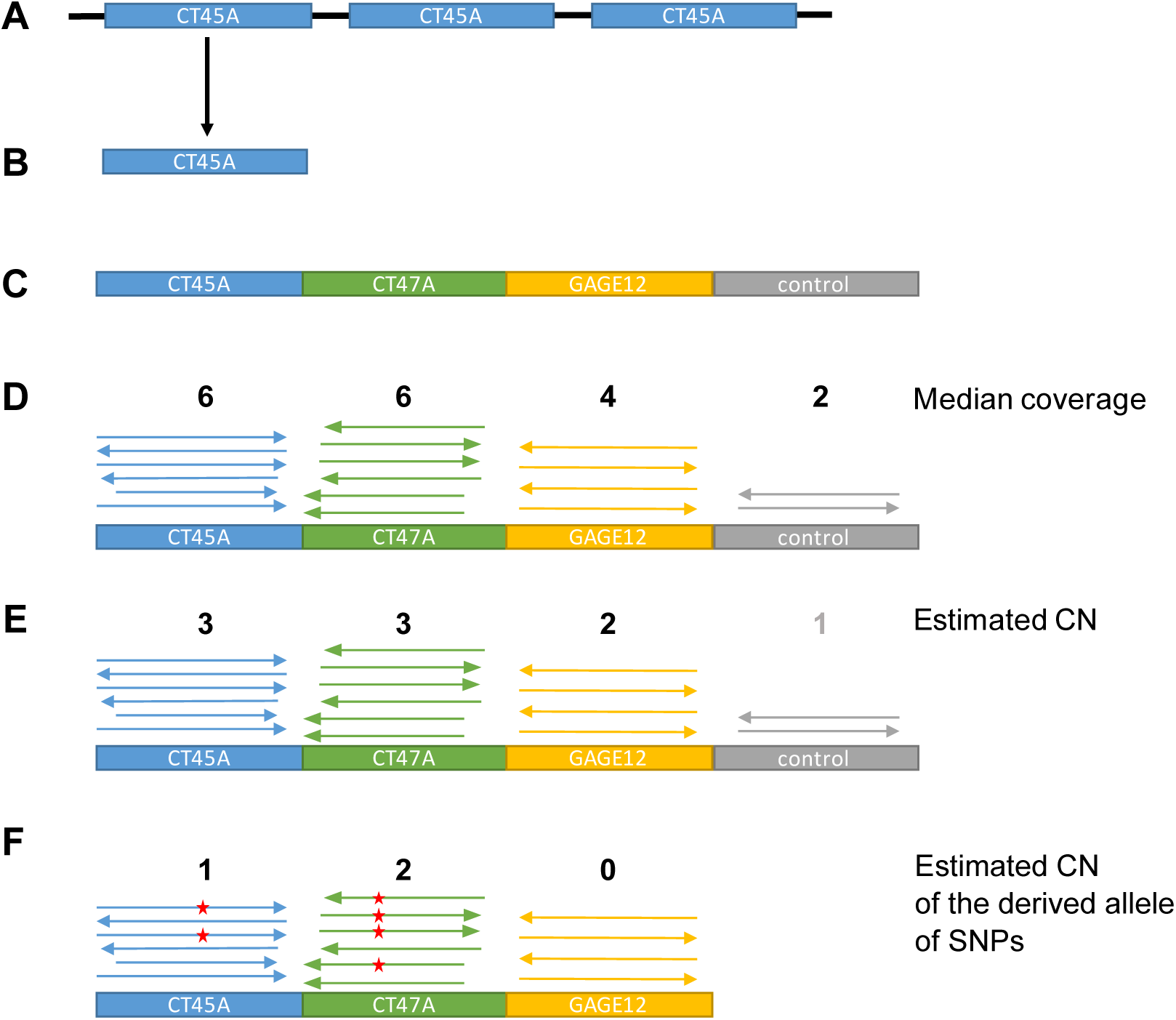
Schema of the artificial X chromosome construction. **A-** Identification of the structure of the ampliconic region using dotplots. **B-** Selection of one unit of repetition of the ampliconic gene. **C-** The sequence of the units of repetition for each ampliconic gene are put next to each other to form an artificial, shorter version of the X chromosome. A control region, known to be single copy, is added. **D-** For each individual, the reads are mapped to the artificial X chromosome, and the median coverage of each unit of repetition and the control region is calculated. **E-** The median coverage of each unit of repetition is divided by the median coverage of the control region to estimate the copy number of each ampliconic genes. **F-** Variant calling is performed on the alignment, and the copy number of the derived allele of each SNP is estimated using coverage.

The coordinates of the X-linked ampliconic regions were taken from Mueller et al. 2013 (Mueller et al. 2013) and were converted to hg38 coordinates. The ampliconic region sequences were extracted from the human reference genome hg38. The sequences of the ampliconic regions were aligned to themselves using *Lastz* (Harris 2007) (--step=100 --notransition --exact=100 --format=rdotplot), and the alignments were filtered to keep only matches longer than 100b. We created dotplots to identify the repeated regions.

The same method was used on non-overlapping 500 kb windows along the X chromosome to identify potential new ampliconic regions and check the existing ones. The boundaries of regions 4, 13, 17, 21,25, 26, 27, 29, 32 (taken from Mueller et al. 2013) were enlarged. Regions 10, 14, 17, 18, 30, 31 and 32 were divided into two units of repetition, and region 29 in three units of repetition. One new region was identified (34), containing the genes CTAG1A, a cancer testis antigen and FAM223A, long intergenic non-protein coding RNA.

We identified the unit of repetition manually on the dotplots. For most of them, the unit was defined by the repeated gene (table S1). For regions that do not contain known genes, we selected the repeated sequence. We identified 24 ampliconic regions on the X chromosome, in which nine are divided into two units of repetition and one is divided into three units of repetition.

We performed a BLAST of the unit sequence against the human reference genome (Altschup et al. 1990). Only one region (14_1) showed 98% of similarity with a region on chromosome 9.

Once the units of repetition were identified, we created mapping targets for raw reads in order to determine the copy number from read depths. We built an artificial chromosome by concatenating the sequences of each unit of repetition plus an X-linked single copy gene, DMD, for control.

#### Y chromosome

On the Y chromosome, the ampliconic regions are structurally more complicated than on the X chromosome, because they are organized in palindromes containing several ampliconic genes (Kuroda-Kawaguchi et al. 2001; Skaletsky et al. 2003). Therefore, we chose to use one copy of each ampliconic genes as units of repetition +− 2kb, as done by Skov and Schierup (2017) (table S2) instead of searching manually for the repeated sequences with alignments. We also included all coding genes from the MSY for controls. An X-degenerate region on the Y chromosome was used as the control region. Therefore, the artificial Y chromosome is composed of 26 genes, including 8 ampliconic genes, and the control region. We also included the sequence of the X chromosome, because most of the Y-linked genes have a closely related X homolog (gametolog). We looked independently at the first two exons of the ampliconic gene PRY because it is known that functional copies of PRY on palindrome 1 do not have the two first exons while copies on palindrome 3 do. For the ampliconic gene RBMY1, we noticed by looking at the coverage of sliding windows that the end of the gene, which does not contain exons, is not always copied. Therefore, we removed this region from the gene sequence.

### Mapping reads against the short chromosomes

To evaluate the number of copy of each ampliconic region, the read files (fastq files from the SGDP dataset) were mapped against the artificial X and Y chromosomes constructed as described above. The read files were also mapped to two control regions: the whole X chromosome excluding the PARs for the X chromosome, and the X-degenerate region for the Y chromosome. The median coverage for each ampliconic region was corrected by the median coverage of the corresponding control region.

*BWA* 0.7.5 (Li and Durbin 2009) was used to perform the mapping (mem -M -t 16 -a). *sambamba* 0.5.1 (Tarasov et al. 2015) was used to filter the paired reads, sort the reads per coordinates, remove the duplicates and filter the bam files for a mapping quality >= 50, an alignment match of at least 100bp and a number of mismatches lower than 2bp. The coverage for each position was obtained using *SAMtools* 1.3 (Li et al. 2009).

### Variant Calling and estimation of the number of copy bearing a variant

A multiple sample variant calling was performed using *platypus* version git-20150612 (Rimmer et al. 2014), without filtering, on males and females for the X chromosome and on males for the Y chromosome. The artificial X and Y chromosomes were used as the reference for the variant calling, so the number of reads supporting a variant will be proportional to the number of copies bearing the variant.

The absence of filtering in the variant calling allowed for the inclusion of variants with allelic imbalance, and the copy number of each variant could be assessed by using the read depth. To estimate the number of copies bearing each variant, we multiplied the estimation of the copy number of the gene for each individual by the number of reads supporting the variant, divided by the number of reads covering the variable position.

Variant calling was also performed on a male chimpanzee (M.H. Schierup, C. Hvilsom, T. Marques Bonet, T. Mailund, unpublished data) to assess the ancestral allele of the variants called in humans. We mapped the fastq files of the chimpanzee to the artificial X and Y chromosomes constructed with the human reference using the same pipeline as for humans. We filtered for 2 mismatches in the alignment and for a length of 100bp. Variant calling was then performed on the alignment using *platypus.* The variant calling in chimp was then confirmed by looking at the base called for each position using the python package *pysam* (http://github.com/pysam-developers/pysam). Both human and chimpanzee vcf files were merged using GATK 3.6 (McKenna et al. 2010) and *picard* 2.7.1 (http://broadinstitute.github.io/picard/). No variants were detected in TSPY.

### Distance trees

Using the R function *NJ* from the package *APE* (Paradis et al. 2004), neighbor-joining trees were constructed on the genetic distances (p-distance) between males for the whole X and the X-degenerate region of the Y chromosome, using SNP data (Nei and Kumar 2000). An alignment was performed on the genotype for each SNP, and a distance matrix was computed for all pair of individuals. The trees were constructed using the R package *ggtree* (Yu et al. 2017).

### Influence of geography and haplogroup on copy number

ANOVA tests were performed on R to assess the influence of the region of origin and haplogroup on copy number. The p-values were corrected using the False Discovery Rate method (Benjamini and Hochberg 1995) for multiple testing over the number of ampliconic regions.

### Signature of selection

The exonic variants were annotated and classified as non-synonymous (aggregating missense variants, stop gained or stop lost variants) or synonymous using *SnpEff* 3.6 (Cingolani et al. 2012) and the annotation database GRCh38.86. A McDonald-Kreitman test and a Direction of Selection test were then performed on each ampliconic regions, and on the pooled ampliconic region to detect signatures of positive selection. Populations were taken into account together and not separately.

### Dataset

We used the Simons Genome Diversity Project, Panel B and C, which includes 274 individuals, 102 females and 172 males, from 128 human populations (Mallick et al. 2016). Panel C samples (260 individuals) were processed using PCR-free library preparation protocol and sequencing protocol. Panel B samples (14 individuals) were processed using a PCR-based library preparation protocol. All samples were submitted to Illumina Ltd. and a 100 base pair paired-end sequencing was performed. The median coverage for the whole sample set we used is of 41.9 with a minimum of 33.59 and a maximum of 83.23 median genome-wide coverage. The median coverage across region varies from 39.35 to 45.

Three individuals were removed from the analysis: *S_Palestinian-2, S_Naxi-2* and *S_Jordanian-1. S_Naxi-2* was removed because it has a high heterozygosity on the X chromosome compared to other males. *S_Palestinian-2* has a low heterozygosity on the X chromosome compared to other females and has an X chromosome coverage comparable to males while having a Y chromosome coverage comparable to female. *S_Jordanian-1* was removed because it is an outlier in all the copy number analysis. These three individuals had also been removed for some analysis in the SGDP paper for different reasons.

## RESULTS AND DISCUSSION

### Copy Number Variations of the Ampliconic Genes

The results for copy number variations of all 34 X-linked regions and 27 Y-linked regions can be found in table S1 and table S2.

Four out of thirty-four X-linked ampliconic regions exhibit extensive copy number variations: Cancer/Testis ampliconic genes CT47A, CT45A, GAGE12 and SPANXB1 (table 1, figure 1). The regions 21_0 and 32_0, as referenced in Mueller et al. (2013), also harbor copy number variation, however, the former does not contain any known gene and the latter contains the gene OPN1LW (opsin 1, long wave sensitive) and we chose to focus on the testis-expressed genes with extensive copy number variation.

**Table 1.** Summary table of the X-linked ampliconic genes with copy number variation

|  | AMPLICONIC REGION |  |  | UNIT OF DUPLICATION |  |  |  | COPY NUMBER VARIATION |  |  |  | ANOVA |
| --- | --- | --- | --- | --- | --- | --- | --- | --- | --- | --- | --- | --- |
| AR | start | end | length | gene | start | end | length | Min CN | Max CN | diff CN | Median CN | qvalue |
| 6_0 | 49219545 | 49680241 | 460696 | GAGE4 | 49560875 | 49568204 | 7329 | 8,84 | 29.35 | 20.51 | 17.23 | **6.70E-04** |
| 24_0 | 120864869 | 120993348 | 128479 | CT47A4 | 120933840 | 120937158 | 3318 | 2.1 | 15.68 | 13.59 | 5.49 | 2.02E-01 |
| 26_0 | 135656801 | 135914069 | 257268 | CT45A5 | 135777129 | 135785475 | 8346 | 3.12 | 10.76 | 7.65 | 7.5 | **6.70E-04** |
| 27_0 | 140964264 | 141635364 | 671100 | SPANXB1 | 140998306 | 141014744 | 16438 | 2.81 | 10.21 | 7.4 | 4.4 | **3.19E-08** |

**Figure 1.**
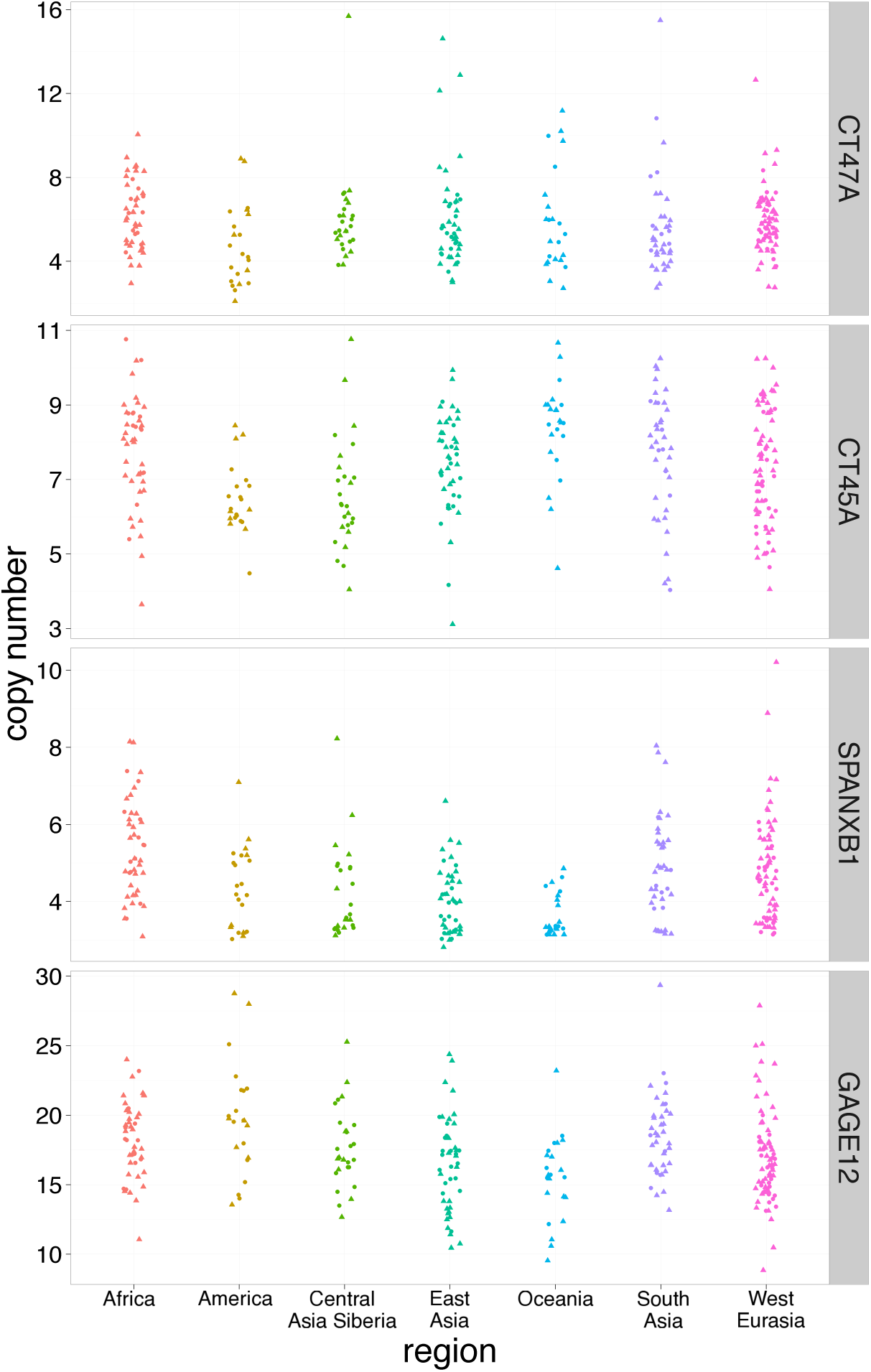
Copy number of X-linked ampliconic genes. Individuals are grouped according to their geographical origin. Females are indicated by circles and males by triangles.

On the Y chromosome, six genes harbor extensive copy number variations: BPY2, CDY, DAZ, PRY, RMBY1A1 and TSPY, all involved in spermatogenesis (table 2, figure 2A-B) while two genes show minor copy number variations: XKRY and HSFY (figure S1). Therefore, all the Y-linked genes defined as ampliconic show copy number variations, except VCY.

**Table 2.**
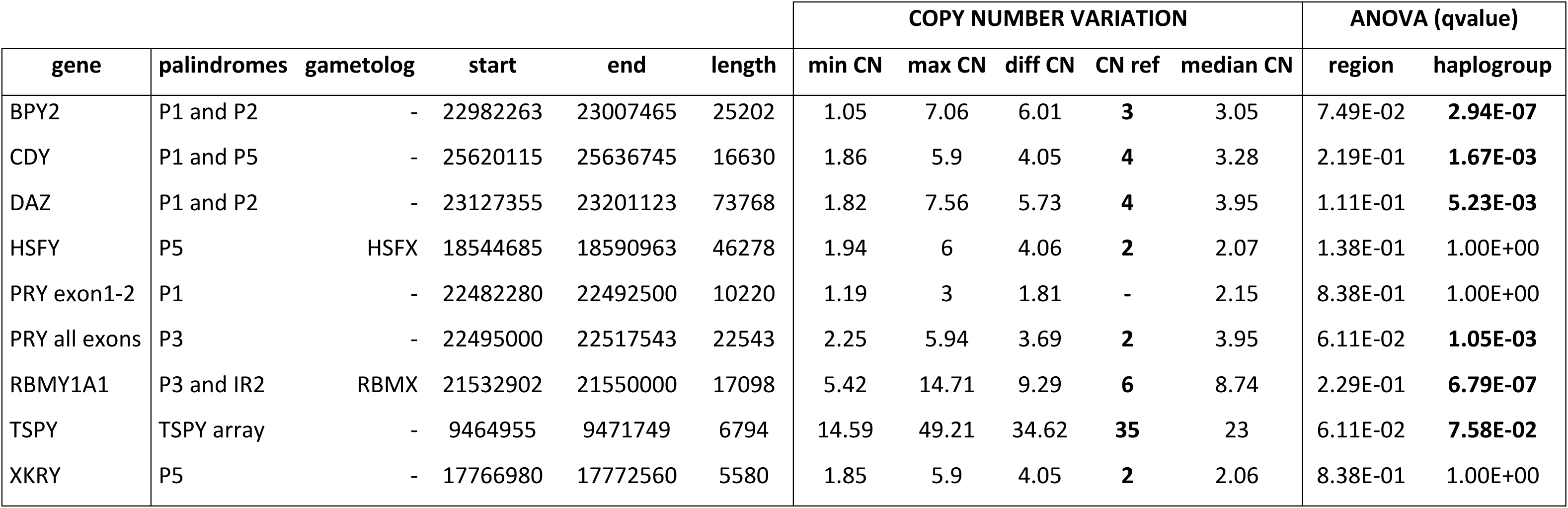
Summary table of the Y-linked ampliconic genes with copy number variation

**Figure 2.**
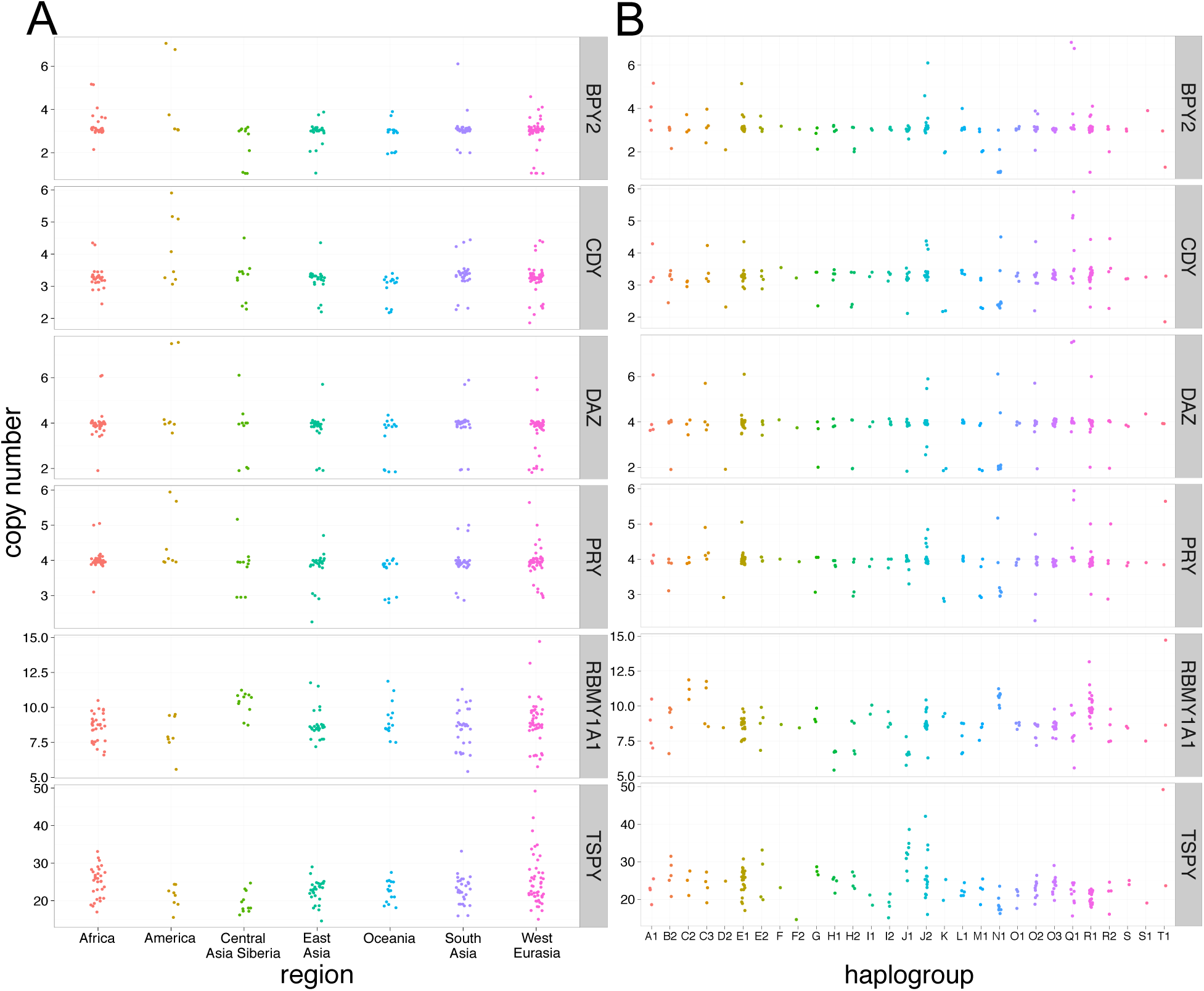
Copy number of Y-linked ampliconic genes. Individuals are grouped according to their **A-** geographical origin and **B-** Y haplogroup.

Henceforth, we focused our analysis on 4 X-linked genes, *i.e.* **CT47A, CT45A, GAGE12 and SPANXB1** and 6 Y-linked genes, *i.e.* **BPY2, CDY, DAZ, PRY, RBMY1A1 and TSPY**.

Our estimation of copy number for each individual is associated with some uncertainty. For Y chromosomes, the copy number can most often be separated into discrete groups, which suggests that our estimates are good. Moreover, a similar method used in Skov et al. (2017) yielded low error rates in father-son pairs. Yet, such discrete classes are not observed for the X chromosome. However, most of the Y ampliconic genes are longer than the X ampliconic genes, which could induce more variance in the coverage. Additionally, recombination events and heterozygous females can further affect the estimation of copy number on X chromosome and not the Y chromosome. As a reference control, the copy number of DMD, an X-linked gene known to be single copy, was calculated and was between 0.95 and 1.17 copies with a median at 1.06 copy per chromosome.

The median copy number for four Y-linked ampliconic genes corresponds to the copy number of the reference Y chromosome (table 2). We find differences for the other ampliconic genes: for PRY we see two additional copies in our study because we accounted for incomplete copies of PRY; for RBMY1A1 we see two additional copies which are most likely pseudogenes not taken into account in the reference; for CDY we see three copies compared to four in the reference, as seen in Skov et al. (2017); and for TSPY the copy number is very different from the reference, probably because this region is highly variable in copy number.

Individuals were separated into geographical groups of origin (figure 1 and 2A). For both X and Y-linked ampliconic genes, copy number variation (CNV) is extensive within geographical groups, the most extreme example on the X chromosome being GAGE12, for which copy number ranges from 9 to 28 in West Eurasia, and the most extreme example on the Y chromosome being TSPY, from 15 to 50 copies in West Eurasia.

Differences in copy number can also be observed between geographical groups. The effect of geographical groups on copy number is significant after correction for multiple testing for 3 out of 4 X-linked genes studied- CT45A, GAGE12 and SPANXB1 (ANOVA, False Discovery Rate (FDR) pvalue<0.05). However, the influence of geography on copy number does not seem to follow any obvious geographical pattern on a map (figure S2).

While none of the Y-linked ampliconic genes show a significant effect of geography on copy number variation (ANOVA FDR pvalue<0.05, figure S3), they all show a significant effect of haplogroups after correction for multiple testing (figure 2B, ANOVA FDR pvalue<0.05). However, copy number variation is repeatedly found within most haplogroups (figure 2B) and distant haplogroups can harbor the same number of copies while closely related haplogroups harbor different copy numbers.

Our results show that amplification and loss of copies happened repeatedly on different branches of the Y haplogroup phylogeny suggesting that changes in copy number occur on a faster scale than the diversification of haplogroups, which mainly occurred within the past 60.000 years (Jobling and Tyler-Smith 2017). This is consistent with previous studies on Y chromosome CNVs (Johansson et al. 2015; Wei et al. 2015; Poznik et al. 2016; Skov et al. 2017).

We also observed large scale variants affecting complete palindromes on the Y chromosome (figure S4). For example, in America, we see that two individuals from the Zapotec population have a duplication of palindrome 1 and 2- *i.e.* 8 copies of DAZ (+4 copies compared to the reference), 7 copies of BPY (+4), 4 copies of PRY_3-4-5_ (+2) and 5 copies of CDY (+2). The individual S_Karitiana-1 has one complete and one incomplete duplication of palindrome 5- *i.e.* 6 HSFY (+4), 6 CDY (+3), 6 XKRY (+4). In West Eurasia, two individuals (S_Adygei-1, S_Tuscan-2) have a duplication of palindrome 2- *i.e.* 6 DAZ (+2), 4 BPY2 (+1).

The correlation between copy number of every possible pair of ampliconic genes on the X and on the Y chromosome was calculated, both with all populations taken together and separately. After correction for multiple testing (False Discovery Rate), no significant correlation was detected (table S3). This suggests that there is not a simple pattern of co-amplification between the X and Y-linked ampliconic genes like seen in mice.

Although we observe extensive variation, the distributions of copy number are similar in all regions, suggesting that although the process of amplification and loss of copies is very dynamic, copy number is kept within a relatively limited range and not subject to a runaway amplification process.

### Coding variation within the ampliconic genes

The ampliconic genes are protein-coding and some of the X-ampliconic genes have previously been reported to be under adaptive evolution in primates (Stevenson et al. 2007; Liu et al. 2008; Zhang and Su 2014). We performed single nucleotide variant calling on the aggregated copies of the ampliconic genes, using the alignment of the reads on the modified X and Y chromosomes. We allowed for allelic imbalance since a given variant may only be polymorphic in some of the copies, but our approach does not allow us to tell which copy the variant is located in. Each variant was annotated as intergenic, synonymous (S), or non-synonymous (NS) (table 4, individual gene results shown in table 3, S1 and S2). We used a chimpanzee sequence as an outgroup to determine the ancestral allele of each variant.

**Table 3.**
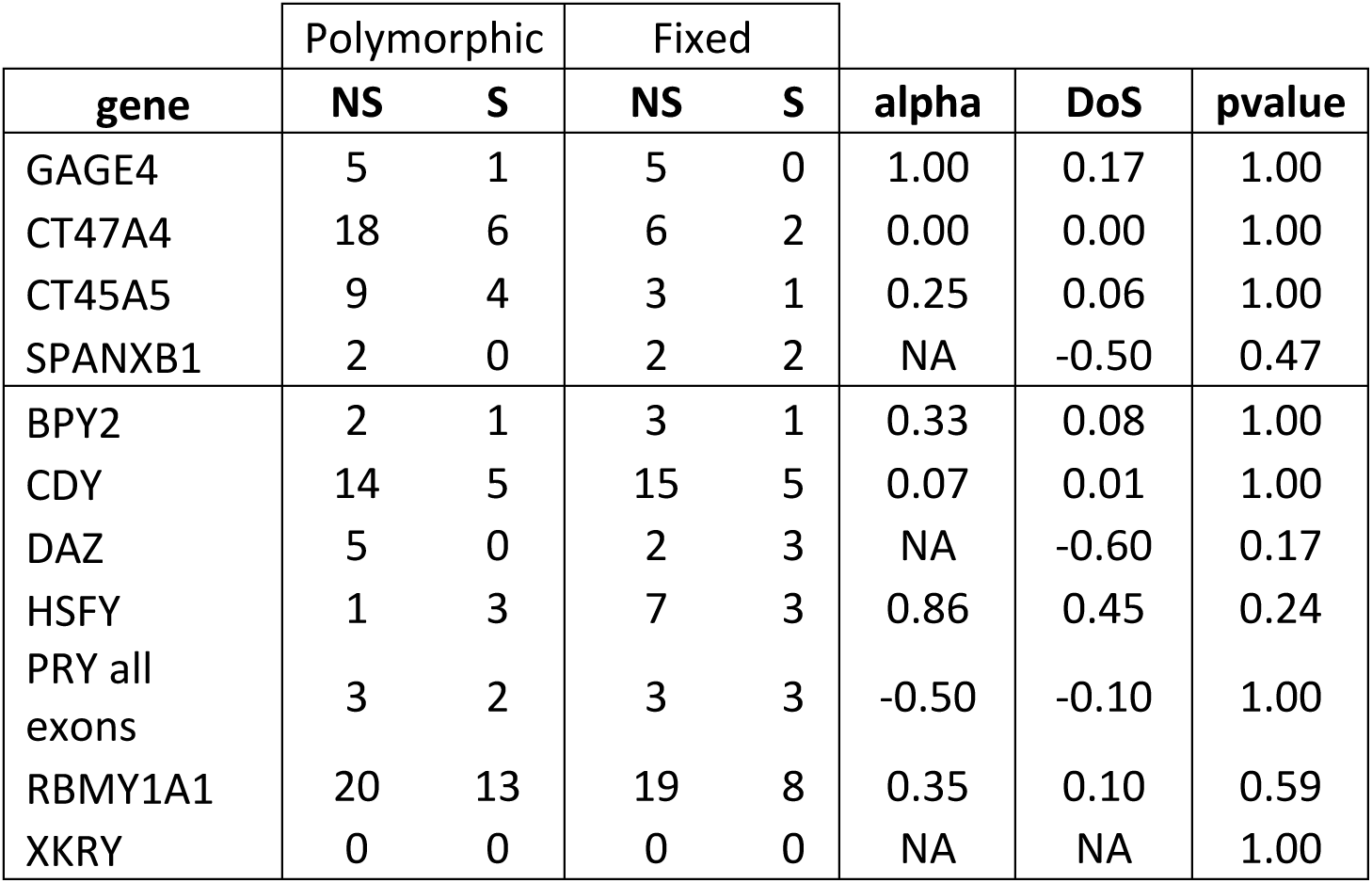
Number of polymorphic and fixed SNPs per gene and results of the Degree of Selection (DoS) and McDonald-Kreitman test (alpha)

**Table 4.** Number of polymorphic and fixed SNPs in the X and Y ampliconic genes

|  | Polymorphic |  | Fixed |  | alpha | DoS |
| --- | --- | --- | --- | --- | --- | --- |
|  | NS | S | NS | S |  |  |
| X (all genes) | 211 | 105 | 102 | 40 | 0.21 | 0.05 |
| Y (all genes) | 95 | 50 | 182 | 102 | -0.06 | -0.01 |

A McDonald-Kreitman test and a Direction of Selection test were performed both on each ampliconic gene and on the joint set of genes, on all populations put together. There is no significant evidence for adaptive evolution (table 3 for the ampliconic genes showing major CNV, table 4 for the pooled genes). Thus, we cannot rule out that these genes are evolving under relaxed purifying selection against amino acid substitutions. However, these selection tests do not take the number of copies of variants into account, which could be crucial as the number of copy might affect the level of expression of the gene.

Next, for the subset of ampliconic genes with ample copy number variations defined above, we investigated the number of copies carrying each derived nonsingleton variant, in relation to the copy number of the gene (see figure S5 for a schematic of the method). We separated common variants, *i.e.* present in at least two copies in at least one individual (figure S6-S9), and rare variants, *i.e.* present in less than two copies (figure S10-S13). We focused on common variants to assess the correlation between the copy number of the gene and the copy number bearing the derived allele of each variant, thus helping us assess the amplification dynamics (table S4).

Several variants show a significant correlation between the copy number bearing the derived allele of the variant and the copy number of the gene: for the X chromosome, nine out of ten NS variants including two with a negative correlation (figure S6), and two out of five S variant with one negative and one positive correlation (figure S7); for the Y chromosome, five out of eight NS variant (figure S8) and two out of three S variants have a significant positive correlation (figure S9).

Due to the multicopy nature of ampliconic genes it is not possible to derive a classical site frequency spectrum for variants. Instead, for the genes selected in this study, we calculated the number of copy bearing the derived allele of each variants in the whole sample and compared it to the summed number of copies of the genes in the whole sample (table 5). For the X chromosome, we find that none of the 15 synonymous variants have a frequency above 10%, whereas this is the case for 10 out of 32 non-synonymous variants. For the Y chromosome, 14 out of 46 NS variants are above 10% frequency, compared to 5 out of 22 NS variants. This suggests that the gene conversion and copy number processes preferentially promote the spread of new non-synonymous variants both among individuals and among gene copies.

**Table 5.** Site frequency spectrum of the NS and S variants

| frequency | X-linked variants |  | Y-linked variants |  |
| --- | --- | --- | --- | --- |
|  | NS | S | NS | S |
| 0 | 32 | 15 | 32 | 17 |
| 0.1 | 3 | 0 | 2 | 2 |
| 0.2 | 0 | 0 | 3 | 1 |
| 0.3 | 2 | 0 | 2 | 2 |
| 0.4 | 0 | 0 | 1 | 0 |
| 0.5 | 0 | 0 | 2 | 0 |
| 0.6 | 0 | 0 | 1 | 0 |
| 0.7 | 1 | 0 | 1 | 0 |
| 0.8 | 2 | 0 | 1 | 0 |
| 0.9 | 2 | 0 | 1 | 0 |

Most of the variants that are present in more than two copies are widespread across populations, with no clear geographical pattern of amplification (figure S6-S9). For example, one NS variant in CT45A (position 135779564) shows a strong correlation between the copy number bearing the derived allele and the gene copy number, except for several African individuals who do not follow this trend (figure 3). Overall, individuals with the same copy number do not necessarily have the same copies duplicated because some do not harbor the variant while other individuals have the variants in several copies.

**Figure 3.**
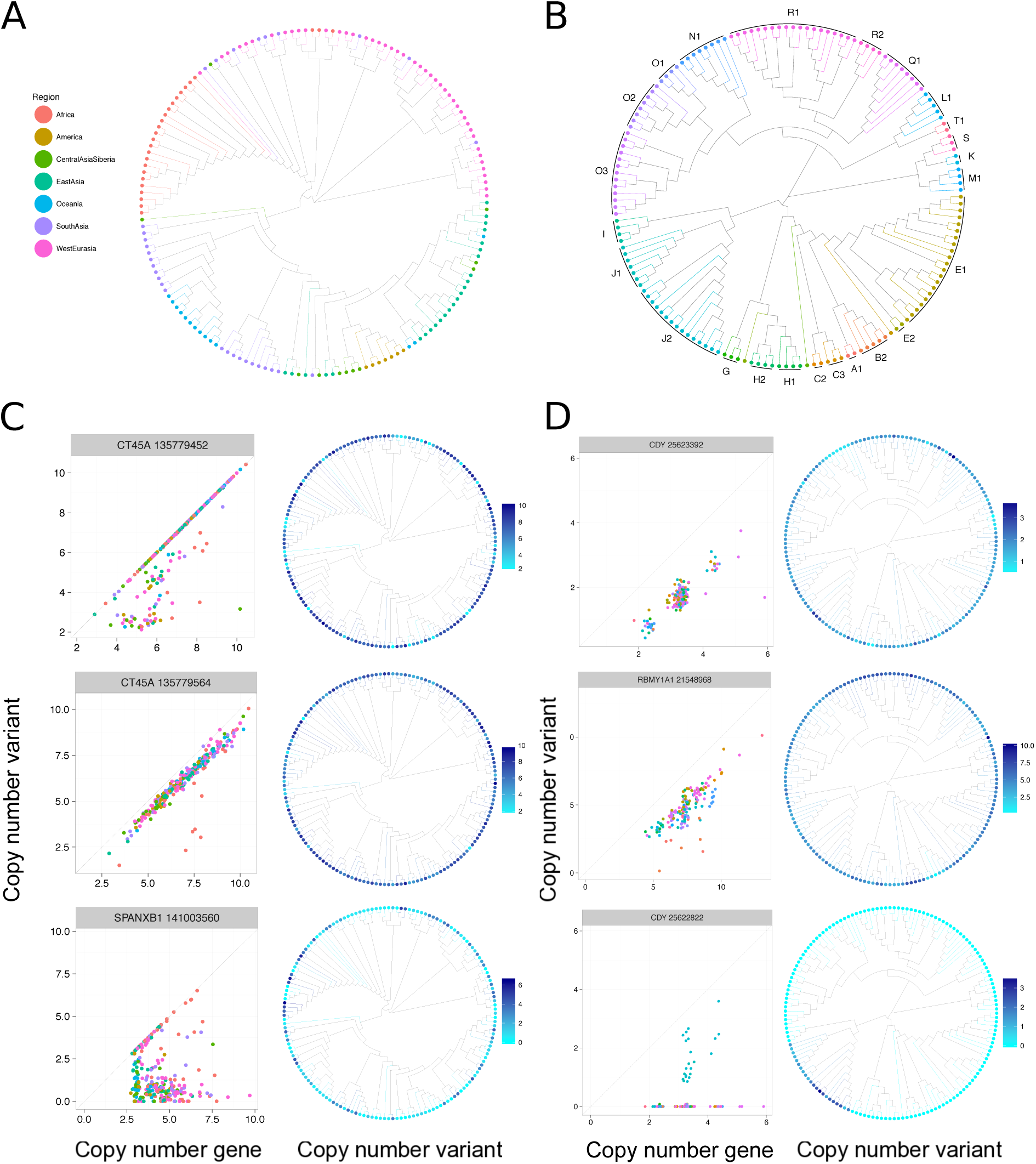
Distribution of copy number bearing the derived allele of variants. Tree of the neighbor joining distance between males for **A-** the X chromosome colored according to the geographical origin of each individual and **B-** the X-degenerate region of the Y chromosome colored according to the haplogroup of each individual. Number of copies of the derived alleles compared to the number of copy of the gene for each individual for three representative example of **C-** X-linked NS variants **D-** Y-linked NS variants. Next to these plots are the distance trees colored according to the copy number of the derived allele of each individual for the variant represented on the left.

The pattern of positive correlation that we observe for many variants can be explained by two mechanisms: duplication of copies bearing the variant or gene conversion. We cannot rule out that gene conversion occurs at high rates. However, if only gene conversion was at play without any duplication, we would only see a vertical relationship between the copy number of the gene and the variant, like seen on figure 3 for the SPANXB1 variant for a number of copies of the gene approximately equal to three. Therefore, amplification of the copies bearing the derived allele happened for most of the variants, especially the NS variants.

To study this pattern and further assess how fast amplification and loss of copies occurred, we constructed neighbour-joining trees with the genetic distance between males for the whole X chromosome and the X-degenerate region of the Y chromosome using SNPs and colored the leaves according to the copy number of each variant present in more than 2 copies (figure S15).

Globally, we can see that individuals that are closely related do not necessarily bear the same copy number of a variant. Here, we focused on 3 representative examples of NS variants for the X and the Y chromosomes, shown in figure 3.

For the Y chromosome, because it does not recombine outside of the pseudoautosomal regions, we can infer multiple events of loss and amplifications of copies bearing some variants (CDY 25623392 and RBMY1A1 21548968) while CDY 25622822 represents a simple case of emergence and amplification of a variant in one branch. Individuals with the same Y haplogroup can bear different copy number of the ampliconic genes and of the derived allele of the variants, which means that amplification and loss happened after the differentiation with their common ancestor, between 60 kya and 30 kya. We can conclude that independent loss and amplification of ampliconic genes have happened and, importantly, that the events happened since the diversification of the Y haplogroups, which suggests that this process is extremely fast.

For the X chromosome, we observe the same pattern as for most of the Y chromosome variants: individuals that are closely related do not have the same copy number of the variant. However, because of recombination, we cannot infer that these events are independent, but it seems that diversity has been kept within populations for these ampliconic regions, both in terms of copy number of genes and variants.

We detected more NS variants than S variants in the ampliconic genes studied here: 69% of the variants are NS for the X ampliconic genes and 70% of the variants are NS for the Y ampliconic genes. We also observe a positive correlation between the variant CN and the gene CN more often for NS variants than for S variants. This observation could be explained by selective forces driving a rapid differentiation of these regions, or by a neutral process that happens faster because of the ampliconic nature of these regions.

We showed above that we do not observe selection signature, but selection tests do not take into account the number of copies bearing the variant. It is possible that the number of copies bearing a variant rather than the variant itself is under selection. The discrepancies we observe between individuals from the same population could be due to complex selective events due to X-Y conflict happening not on copy number, but on combinations of copy number and allele-matching processes. High diversity seems to have been kept within populations, which might suggest balancing selection on both Y and X chromosome ampliconic genes.

### Expression during meiosis

The sex chromosomes are inactivated at the end of meiosis, during pachytene and diplotene (MSCI), and remain repressed during spermiogenesis (PSCR). However, some genes can escape this process and still be expressed during these stages (Sin et al. 2012). It has been suggested that amplification allows genes with an important function in spermatogenesis to increase their expression and therefore counterbalance the repressive effect of MSCI and PSCR.

Using the data from Sin et al. (2012), we looked at the expression of the ampliconic genes showing major CNV on the X chromosome (CT45A, CT47A, GAGE1 and SPANXB1 gene families, figure 4A) and on the Y chromosome (BPY2, CDY1, DAZ, HSFY, PRY, RBMY1, TSPY and XKRY gene families, figure 4B) in humans. These data inform us about the level of expression in three cell types with increasing differentiation during spermatogenesis: spermatogonia (SG) before meiosis, pachytene spermatocytes (PS) during meiosis and MSCI, and round spermatids (RS) during post-meiotic sex chromosome repression (PSCR).

**Figure 4.**
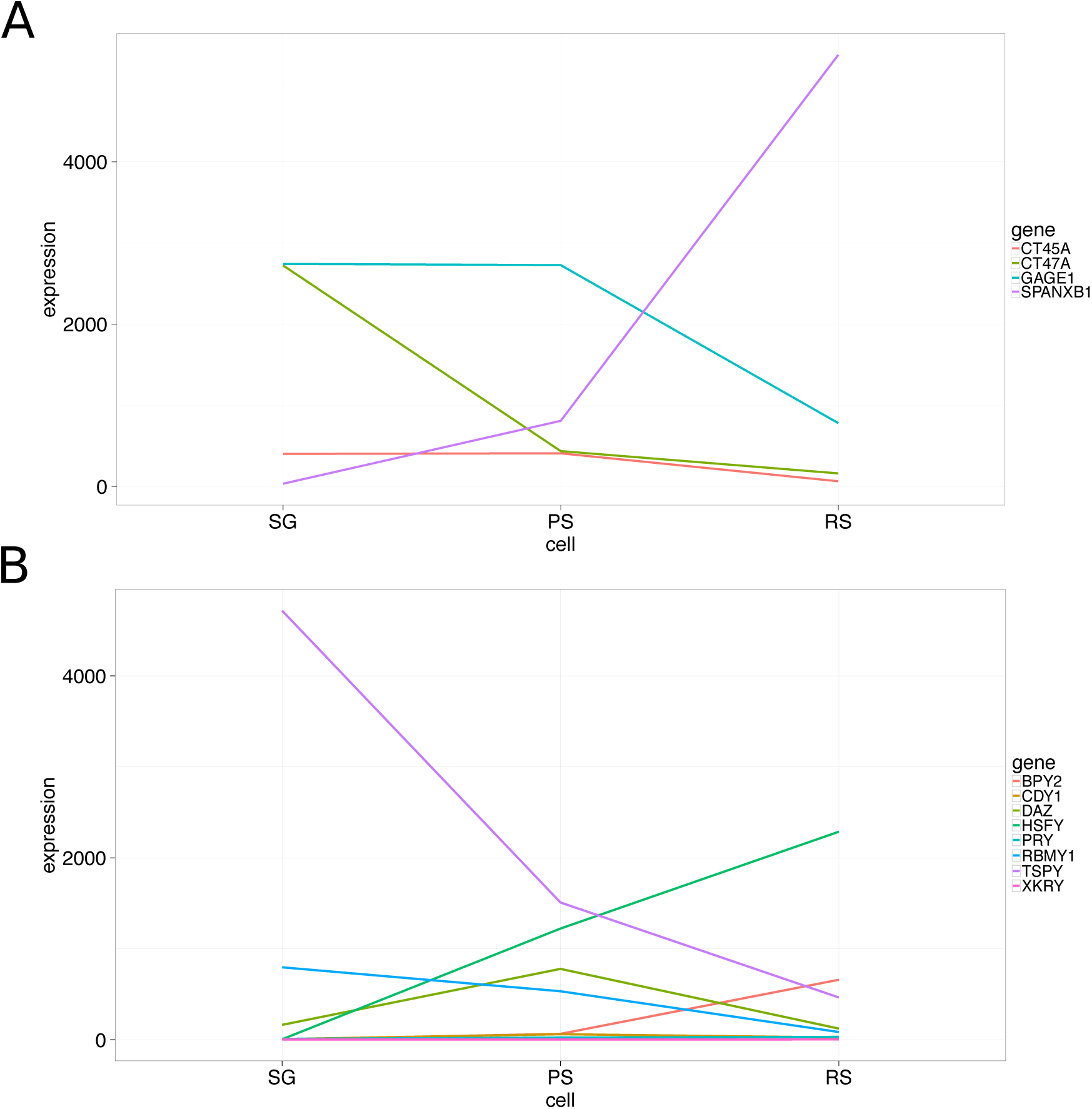
**Levels of expression during different stages of male meiosis** of **A-** X-linked ampliconic genes and **B-** Y-linked ampliconic genes. On the X axis, different cell types are represented: spermatogonia (SG) before meiosis, pachytene spermatocytes (PS) during meiosis and meiotic sex chromosome inactivation, and round spermatids (RS) during post-meiotic sex chromosome repression. Expression data were taken from Sin et al 2012.

We find that the X-linked genes showing high copy number variations are all expressed in pachytene spermatocytes during MSCI, and two of them are expressed during PCSR (SPANXB1 and GAGE1). The SPANXB1 gene family is the most striking example, with an expression that increases through meiosis and more dramatically during PSCR. Four of the Y-linked ampliconic genes showing high CNV are expressed during MSCI in pachytene spermatocytes (TSPY, HSFY, DAZ and RBMY1 families), and three gene families are expressed in round spermatids during PSCR (TSPY, BPY2 and HSFY). Moreover, BPY2 is expressed only during PSCR, which indicates that it is inactivated during MSCI and reactivated during PSCR.

Sin et al. (2012) showed that escape genes in humans are often de novo genes that appeared in the primate or great ape lineage and that they show a higher Ka/Ks ratio than non-escape genes, and therefore undergo a faster rate of evolution. This is concordant with the dynamic copy number gain/loss and the high mutation rate of the ampliconic genes highlighted in our study.

Our genes of interest are expressed specifically in testis during meiosis and are good candidate genes for hybrids incompatibilities emergence. Interestingly, in humans, regions depleted of Neanderthal and Denisovan ancestry are enriched for genes expressed in the testis (Sankararaman et al. 2016) and particularly in genes expressed during meiosis (Jégou et al. 2017).

Further studies on the impact of copy number variations and copy number of variants on gene expression and sperm phenotypes will allow us to assess the role of the ampliconic genes in hybrid incompatibility emergence.

## CONCLUSION

We have found that copy number variation is extensive within and between populations for four X-linked genes and six Y-linked genes. Although geography (for the X) and haplogroup (for the Y) can explain some of the variability, we do not find a clear geographical pattern, and individuals closely related can harbour very different copy number and variants. Thus, our results demonstrate an extremely fast copy number variation dynamic of Y-linked ampliconic genes. For the X chromosome, because of recombination, we cannot conclude that the amplification events observed are independent. However, we can say that diversity seems to have been kept within populations, which suggests that maintenance of diversity in these regions is important. We propose that complex selective events might have occurred on the copy number of variants, in an allele-matching process between Y and X ampliconic genes. To disentangle neutral processes and selection on copy number, one would need to construct a null model, *i.e.* what would be the diversity patterns if only drift was at play, while taking into account the many peculiarities of these regions and gene conversion. This study is a first step toward this goal.

The ampliconic genes highlighted here are expressed in testis and during stages of meiosis where the sex chromosomes are inactivated and knowing that genes involved in meiosis are depleted of Neanderthal ancestry, it suggests that ampliconic genes could be linked to hybrid incompatibilities.

While we cannot yet disentangle neutral processes and selective forces acting on copy number, we argue that it is unlikely that copy number variation on such important genes, that are expressed during spermatogenesis, are completely neutral. Moreover, while we see important copy number variation between individuals, the copy number seems to be constrained and not under a runaway process, which suggests that having too many or too few copies is deleterious. Furthermore, only non-synonymous variations are observed in high frequency among copies, suggesting positive selection might be at play.

Further studies on the co-evolution of these regions between sex chromosomes across primate species, and on the functional impact of copy number variations on spermatogenesis have been initiated.

## ACKNOWLEDGEMENTS

The authors would like to thank members of the Bioinformatics Research Center, especially Jacob Malte Jensen, David Castellano and Thomas Bataillon for discussions on the manuscript. This work was supported by funding from the Danish Research Council for Independent Research.

## REFERENCES

Altschup SF, Gish W, Miller W, Myers EW, Lipman DJ. 1990. Basic Local Alignment Search Tool. J. Mol. Biol. 215:403–410.

Cingolani P, Platts A, Wang LL, Coon M, Nguyen T, Wang L, Land SJ, Lu X, Ruden DM. 2012. A program for annotating and predicting the effects of single nucleotide polymorphisms, SnpEff. Fly (Austin). 6:80–92.

Cocquet J, Ellis PJI, Mahadevaiah SK, Affara NA, Vaiman D, Burgoyne PS. 2012. A Genetic Basis for a Postmeiotic X Versus Y Chromosome Intragenomic Conflict in the Mouse. PLoS Genet. 8.

Davis BW, Seabury CM, Brashear WA, Li G, Roelke-Parker M, Murphy WJ. 2015. Mechanisms underlying mammalian hybrid sterility in two feline interspecies models. Mol. Biol. Evol. 32:2534–2546.

Dutheil JY, Munch K, Nam K, Mailund T, Schierup MH. 2015. Strong Selective Sweeps on the X Chromosome in the Human-Chimpanzee Ancestor Explain Its Low Divergence. PLoS Genet. 11:1–18.

Frank SA. 1991. Divergence of Meiotic Drive-Suppression Systems as an Explanation for Sex-Biased Hybrid Sterility and Inviability. 45:262–267.

Ghenu A-H, Bolker BM, Melnick DJ, Evans BJ. 2016. Multicopy gene family evolution on primate Y chromosomes. BMC Genomics 17:157.

Gjerstorff MF, Ditzel HJ. 2008. An overview of the GAGE cancer/testis antigen family with the inclusion of newly identified members. Tissue Antigens 71:187–192.

Harris RS. 2007. Improved pairwise alignment of genomic DNA.

Hurst LD, Pomiankowski A. 1991. Causes of sex ratio bias may account for unisexual sterility in hybrids: A new explanation of Haldane’s rule and related phenomena. Genetics 128:841–858.

Jégou B, Sankararaman S, Rolland AD, Reich D, Chalmel F, Affiliations ¤. 2017. Meiotic genes are enriched in regions of reduced archaic ancestry. Mol. Biol. Evol. 21:1974–1980.

Jobling MA, Tyler-Smith C. 2017. Human Y-chromosome variation in the genome-sequencing era. Nat. Rev. Genet. 18:485–497.

Johansson MM, Van Geystelen A, Larmuseau MHD, Djurovic S, Andreassen OA, Agartz I, Jazin E. 2015. Microarray analysis of copy number variants on the human y chromosome reveals novel and frequent duplications overrepresented in specific haplogroups. PLoS One 10:1–23.

Kuroda-Kawaguchi T, Skaletsky H, Brown LG, Minx PJ, Cordum HS, Waterston RH, Wilson RK, Silber S, Oates R, Rozen S, et al. 2001. The AZFc region of the Y chromosome features massive palindromes and uniform recurrent deletions in infertile men. Nat. Genet. 29:279–286.

Larson EL, Keeble S, Vanderpool D, Dean MD, Good JM. 2016. The composite regulatory basis of the large X-effect in mouse speciation. Mol. Biol. Evol. 34:msw243.

Li H, Durbin R. 2009. Fast and accurate short read alignment with Burrows-Wheeler transform. Bioinformatics 25:1754–1760.

Li H, Handsaker B, Wysoker A, Fennell T, Ruan J, Homer N, Marth G, Abecasis G, Durbin R. 2009. The Sequence Alignment/Map format and SAMtools. Bioinformatics 25:2078–2079.

Liu Y, Zhu Q, Zhu N. 2008. Recent duplication and positive selection of the GAGE gene family. Genetica 133:31–35.

Mallick S, Li H, Lipson M, Mathieson I, Gymrek M, Racimo F, Zhao M, Chennagiri N, Nordenfelt S, Tandon A, et al. 2016. The Simons Genome Diversity Project: 300 genomes from 142 diverse populations. Nature 538:201–206.

McKenna A, Hanna M, Banks E, Sivachenko A, Cibulskis K, Kernytsky A, Garimella K, Altshuler D, Gabriel S, Daly M, et al. 2010. The Genome Analysis Toolkit: A MapReduce framework for analyzing next-generation DNA sequencing data. Genome Res. 20:1297–1303.

Mueller JL, Mahadevaiah SK, Park PJ, Warburton PE, Page DC, Turner JMA. 2008. The mouse X chromosome is enriched for multicopy testis genes showing postmeiotic expression. Nat. Genet. 40:794–799.

Mueller JL, Skaletsky H, Brown LG, Zaghlul S, Rock S, Graves T, Auger K, Warren WC, Wilson RK, Page DC. 2013. Independent specialization of the human and mouse X chromosomes for the male germ line. Nat. Genet. 45:1083–1087.

Nam K, Munch K, Hobolth A, Dutheil JY, Veeramah KR, Woerner AE, Hammer MF, Mailund T, Schierup MH. 2015. Extreme selective sweeps independently targeted the X chromosomes of the great apes. Proc. Natl. Acad. Sci. 112:6413–6418.

Nei M, Kumar S. 2000. Molecular Evolution and Phylogenetics. New York: Oxford University Press

Oetjens MT, Shen F, Emery SB, Zou Z, Kidd JM. 2016. Y-Chromosome Structural Diversity in the Bonobo and Chimpanzee Lineages. Genome Biol. Evol. 8:2231–2240.

Paradis E, Claude J, Strimmer K. 2004. APE: Analyses of phylogenetics and evolution in R language. Bioinformatics 20:289–290.

Poznik GD, Xue Y, Mendez FL, Willems TF, Massaia A, Wilson Sayres MA, Ayub Q, McCarthy SA, Narechania A, Kashin S, et al. 2016. Punctuated bursts in human male demography inferred from 1,244 worldwide Y-chromosome sequences. Nat. Genet. 48:593–599.

Rimmer A, Phan H, Mathieson I, Iqbal Z, Twigg SRF, Wilkie AOM, McVean G, Lunter G. 2014. Integrating mapping-, assembly- and haplotype-based approaches for calling variants in clinical sequencing applications. Nat. Genet. 46:1–9.

Sankararaman S, Mallick S, Patterson N, Reich D. 2016. The Combined Landscape of Denisovan and Neanderthal Ancestry in Present-Day Humans. Curr. Biol. 26:1241–1247.

Simpson AJG, Caballero OL, Jungbluth A, Chen Y-T, Old LJ. 2005. Cancer/testis antigens, gametogenesis and cancer. Nat. Rev. Cancer 5:615–625.

Sin H, Ichijima Y, Koh E. 2012. Human postmeiotic sex chromatin and its impact on sex chromosome evolution. Genome Res.:827–836.

Skaletsky H, Kuroda-Kawaguchi T, Minx PJ, Cordum HS, Hillier L, Brown LG, Repping S, Pyntikova T, Ali J, Bieri T, et al. 2003. The male-specific region of the human Y chromosome is a mosaic of discrete sequence classes. Nature 423:825–837.

Skov L, The Danish Pan Genome Consortium, Schierup MH. 2017. Analysis of 62 hybrid assembled human Y chromosomes exposes rapid structural changes and high rates of gene conversion. PLOS Genet. 13:e1006834.

Soh YQS, Alföldi J, Pyntikova T, Brown LG, Graves T, Minx PJ, Fulton RS, Kremitzki C, Koutseva N, Mueller JL, et al. 2014. Sequencing the mouse y chromosome reveals convergent gene acquisition and amplification on both sex chromosomes. Cell 159:800–813.

Stevenson BJ, Iseli C, Panji S, Zahn-Zabal M, Hide W, Old LJ, Simpson AJ, Jongeneel CV. 2007. Rapid evolution of cancer/testis genes on the X chromosome. BMC Genomics 8:129.

Tarasov A, Vilella AJ, Cuppen E, Nijman IJ, Prins P. 2015. Sambamba: Fast processing of NGS alignment formats. Bioinformatics 31:2032–2034.

Wei W, Fitzgerald T, Ayub Q, Massaia A, Smith BB, Dominiczak AA, Morris AA, Porteous DD, Hurles ME, Tyler-Smith C, et al. 2015. Copy number variation in the human Y chromosome in the UK population. Hum. Genet. 134:789–800.

Yu G, Smith DK, Zhu H, Guan Y, Lam TTY. 2017. Ggtree: an R Package for Visualization and Annotation of Phylogenetic Trees With Their Covariates and Other Associated Data. Methods Ecol. Evol. 8:28–36.

Zhang Q, Su B. 2014. Evolutionary origin and human-specific expansion of a Cancer/testis antigen gene family. Mol. Biol. Evol. 31:2365–2375.

Zhao Q, Caballero OL, Simpson AJG, Strausberg RL. 2012. Differential Evolution of MAGE Genes Based on Expression Pattern and Selection Pressure. PLoS One 7.

